# KDM: embedding DNA/RNA motifs and sequences in a shared k-mer space for unified discovery, analysis and binding prediction

**DOI:** 10.64898/2026.06.05.730329

**Authors:** Luca Fumagalli, Tommaso Becchi, Matteo Cereda, Uberto Pozzoli

## Abstract

Motif discovery and binding-site prediction in DNA and RNA sequences are central tasks in regulatory genomics, yet the methodological landscape is split between interpretable but rigid position weight matrices (PWMs) and high-performing but opaque machine-learning models. We present KDM, a unifying framework in which both motifs and sequences are represented as probability distributions over a shared k-mer dictionary, embedded via the Hellinger transformation. This common geometry enables motif-sequence scoring, motif-motif comparison, de novo discovery, and binding prediction with a single primitive, the Bhattacharyya coefficient. We instantiate four tools on this representation: KDMMap for positional enrichment analysis, KDMMatch for information-content-aware motif matching, KDMFind for unsupervised motif discovery via projective non-negative matrix factorization, and KDM-LRLM for binding prediction with Lasso-regularized logistic regression. Across 1,324 transcription-factor ChIP-seq and 161 RBP eCLIP experiments, KDMMap matches CentriMo’s motif rankings in 84% of TF and 79% of RBP experiments, and KDMMatch agrees with Tomtom on motif annotation in 74.5% of TFs. On binding prediction across four datasets covering 2,475 experiments, KDM-LRLM matches or exceeds eight deep-learning and three k-mer-based competitors. Notably, AI methods overtake k-mer methods only in the top quartile of training-set size, indicating that data scale, not architecture, drives the recent dominance of deep models. KDM provides a single interpretable representation across the full motif-analysis workflow.

## Introduction

Motif discovery from sequence aims to identify recurring patterns in DNA or RNA that underlie specific protein–nucleic acid interactions, such as transcription factor binding to promoter regions or RNA-binding proteins recognizing exons or introns. These motifs provide a representation of binding specificity and help explain why certain genomic regions are bound in biological experiments. By inferring motifs directly from sets of bound versus unbound sequences, computational methods highlight the core nucleotides and flanking context that are most informative for binding. Ultimately, motif discovery bridges raw sequence data with mechanistic models of gene regulation and post-transcriptional control.

A large class of motif discovery methods relies on position matrices to represent binding preferences^1,2^ (**Supplementary Table S1**). In such matrices, each position is assigned a probability, or log-odds weight, for each nucleotide, assuming that contributions of different positions are independent. We refer to these matrices collectively as Positional Weight Matrices (PWMs). Classical algorithms such as expectation-maximization and Gibbs sampling iteratively refine the PWM by maximizing the likelihood that observed bound sequences were generated from motif and background models. PWM-based approaches are attractive because they are simple, interpretable, and can be visualized as sequence logos. They integrate naturally into downstream analyses, including comparison to annotated motif collections, identification of enriched genomic regions, genome scanning for putative binding sites, and prediction of point-mutation effects (**Fig.1A**).

**Figure 1.**
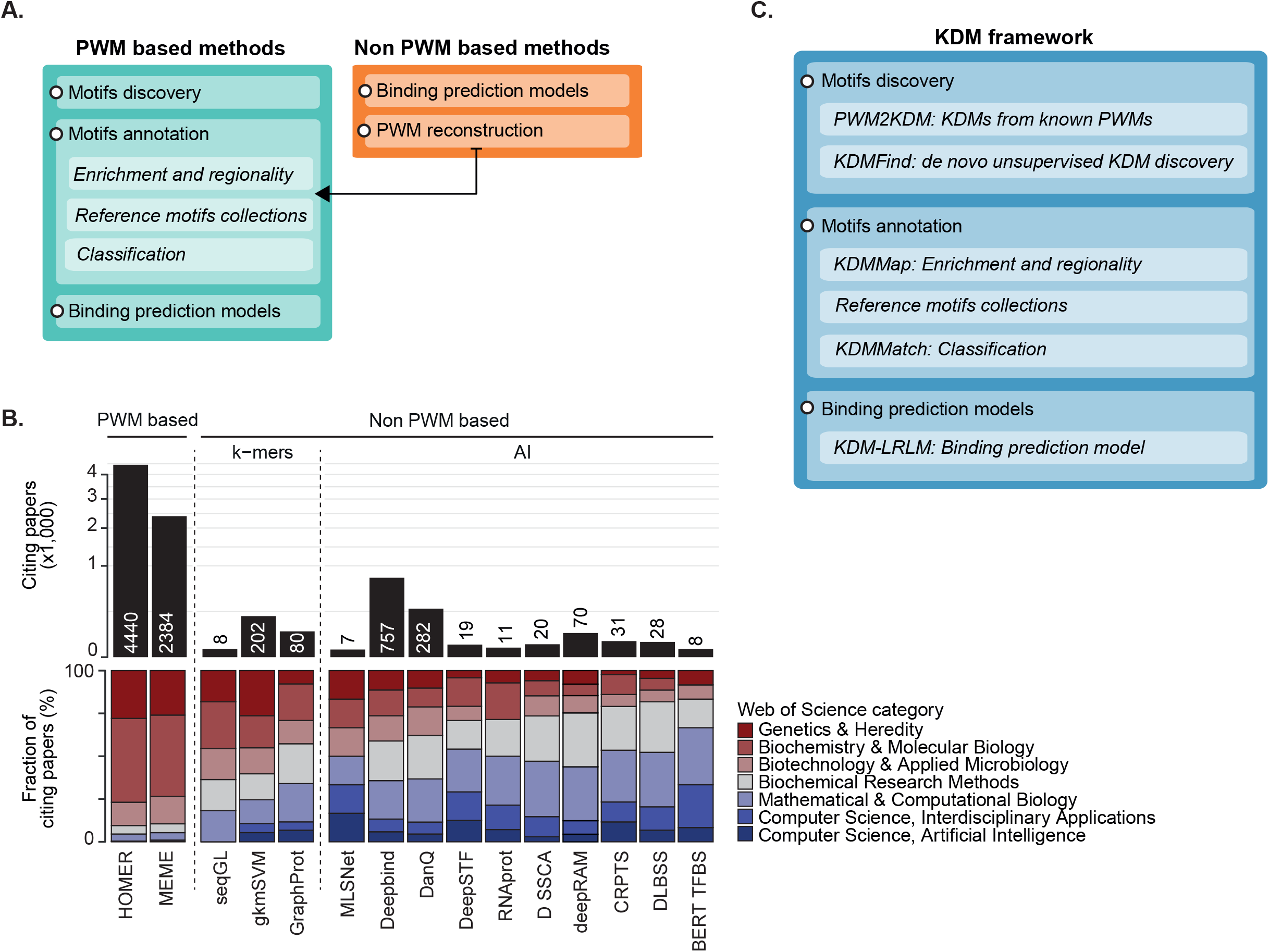
Overview of motif analysis approaches and the KDM framework. **A**. Schematic of key characteristics of PWM-based vs. non-PWM motif analysis approaches. **B**. Publication counts citing each method over the last five years, retrieved from Scopus (Elsevier) from 2021-01-01 to 2025-11-05. For each method, citing publications were stratified by Web of Science category and shown as proportional contributions (bottom; see Methods). **C**. Schematic of the main components of the KDM framework, mapping each tool to the corresponding PWM-based counterpart.

However, the independence assumption built into PWMs often fails to capture the full complexity of protein-nucleic acid recognition. Many proteins exhibit positional dependencies, spacing constraints, or require particular combinations of bases that are not well modeled by additive, position-wise contributions^3^. To address these limitations, a growing set of non-PWM methods has been developed. These include machine-learning approaches such as support vector machines, generalized linear models, and graph-based encodings, typically built on k-mer enrichment ^4–7^ (**Supplementary Table S1**). Such models can capture dependencies across positions or variable-length binding patterns, including motifs with gaps, repeats, or secondary-structure preferences—features especially relevant for RNA-binding proteins. More recently, deep-learning approaches have expanded the non-PWM toolkit, learning nonlinear relationships between sequence features and binding and inferring complex motifs and their combinatorial interactions directly from raw sequence ^8–17^.

Non-PWM models often achieve strong predictive performance and can implicitly capture both sequence motifs and higher-order context such as motif spacing or orientation. The trade-off is reduced transparency. Interpreting learned filters or feature importance typically requires specialized tools, and most k-mer and AI-based methods ultimately reconstruct PWMs from learned parameters for downstream visualization and annotation. Together, PWM-based and non-PWM methods form a complementary spectrum of approaches for dissecting the sequence determinants of protein-DNA/RNA interactions. Yet, these approaches are still largely applied in isolation, often for different purposes. Researchers with a biological background tend to rely on traditional PWM-based approaches because of their greater interpretability and ease of use, creating a persistent disconnect between the rapid development of high-performing computational methods and their effective adoption in biological research (**Fig. 1B**).

Here, we present KDM, a k-mer-based framework that represents motifs as probability distributions over a shared k-mer dictionary (K-mer Distribution Motifs, KDMs). The central insight is that, under the Hellinger embedding, both motifs and sequences occupy the same geometric space, so a single primitive, the Bhattacharyya coefficient, suffices to (*i*) score motifs in sequences, (*ii*) compare motifs to each other, and (*iii*) drive unsupervised motif discovery. KDMs can be derived analytically from existing PWMs or discovered *de novo* through Projective Non-negative Matrix Factorization (PNMF). Building on this representation, we develop four tools that together cover the standard motif-analysis workflow: (*i*) KDMMap for positional enrichment, (*ii*) KDMMatch for information-content (IC) aware annotation, (*iii*) KDMFind for de novo discovery, and (*iv*) KDM-LRLM for binding prediction (**Fig. 1C**). We benchmark each tool against established PWM and non-PWM counterparts on large-scale TF and RBP datasets, and report a counter-intuitive finding: across more than 2,000 binding-prediction experiments, deep-learning methods overtake k-mer methods only when training-set size is large, suggesting that data scale rather than model architecture explains much of the recent dominance of AI approaches.

## Results

Our framework represents sequence-motif patterns as *probability* distributions over k-mers, referred hereafter as k-mer distribution motifs (KDMs), and encodes nucleotide sequences as vectors of observed k-mer *frequencies* in the same dictionary, enabling direct comparison and scoring between motifs and sequences.

Below, we first define the KDM representation, sequence encoding, and similarity measures, then build on these definitions to generalize widely used PWM-based analytical operations to KDMs and extend them to predictive modeling (**Fig. 1A-C**, as described above).

### KDM representation and encoding

We represent both motifs and sequences in the same k-mer feature space defined by a dictionary *D*. A motif is encoded as a probability distribution over k-mers, a KDM, while a sequence is the empirical distribution of k-mer frequencies observed within it. This unified representation enables direct comparison between motifs and sequences, and between motifs themselves, using standard measures of distributional similarity. We define below the dictionary, the KDM representation, the sequence encoding, and the similarity measures used throughout.

#### K-mer dictionary *D*

We define *D* as the set of DNA/RNA words of length k (k-mers), optionally including the symbol N to indicate gaps. *D* can be double-stranded, where a k-mer and its reverse complement are treated as the same dictionary entry, or single-stranded, where all k-mers are distinct. For DNA, we used a double-stranded dictionary with 8-mers (k = 8) allowing two gaps, comprising 57,472 unique entries. For RNA, we exploit a single-stranded dictionary with 6-mers (k = 6) allowing two gaps, comprising 3,840 k-mers. The DNA parameters follow established choices in gapped k-mer methods (gkmSVM^18^, SeqGL^6^); the RNA parameters were selected by grid search (Methods, **Supplementary Fig. S1**)

Given *D*, the core assumptions were (*i*) to represent a motif pattern as a probability distribution p over all k-mers in *D*, where each *p*_*i*_ is the probability that the motif emits k-mer *i*, and (*ii*) to encode a sequence as an empirical distribution *q*, where each *q*_*i*_ is the relative frequency of k-mer *i* observed in that sequence.

To compare two distributions *p* and *q*, we use the Bhattacharyya coefficient (*BC*)^19^, defined for discrete distributions as:

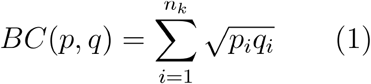

Each term *√(p*_*i*_*q*_*i*_) is the geometric mean of the motif probability and the observed frequency for k-mer *i*. It is non-zero only for k-mers that are both probable under the motif and present in the sequence, and BC sums these contributions across the dictionary. A closely related distance is the Hellinger distance (*HD*) ^20^:

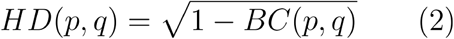

which converts the similarity into a distance: H is largest (HD=1) when the motif and sequence share no k-mers, and smallest (HD=0) when their distributions are identical.

Because motifs (probabilities, *p*) and sequences (frequencies, *q*) are both distributions over the same dictionary, *BC* and *HD* provide a natural way to score how well a motif matches a sequence, and to compare motifs (or sequences) to one another. For compact notation, we worked with square-root-transformed distributions, the Hellinger embedding of a categorical distribution on *D*:

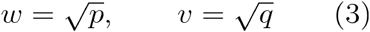

Defining *w* for a motif and *v* for a sequence, the BC reduces to a simple dot product between the two embedded vectors:

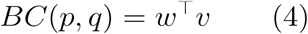

so that scoring a motif against a sequence becomes the geometric overlap (cosine-like inner product) of their square-root distributions.

#### KDM

Given a dictionary *D* of *n* k-mers, a KDM is a motif pattern represented as the vector *w* of square roots of k-mer probabilities over *D* (**Fig.2A**); that is, each entry *w*_*i*_ is the Hellinger embedding of the motif probability *p*_*i*_ for k-mer *i*:

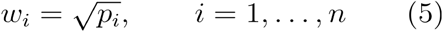

A collection of *n*_*x*_ motifs is represented as a matrix *W*:

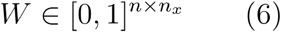

A KDM can also be derived from a PWM using the PWM2KDM procedure (**Fig. 2A**). Given a PWM, we estimate the probability of each k-mer in D by sliding the k-mer across PWM positions. For each alignment, the k-mer probability is obtained by multiplying the PWM probabilities of the aligned letters (positions outside the PWM are treated as uniform). If D is double-stranded, we repeat the calculation with the reverse-complement PWM and average the two results. The final k-mer probability is the average across all alignments, and the KDM entry *w*_*i*_ is obtained as in (5).

**Figure 2.**
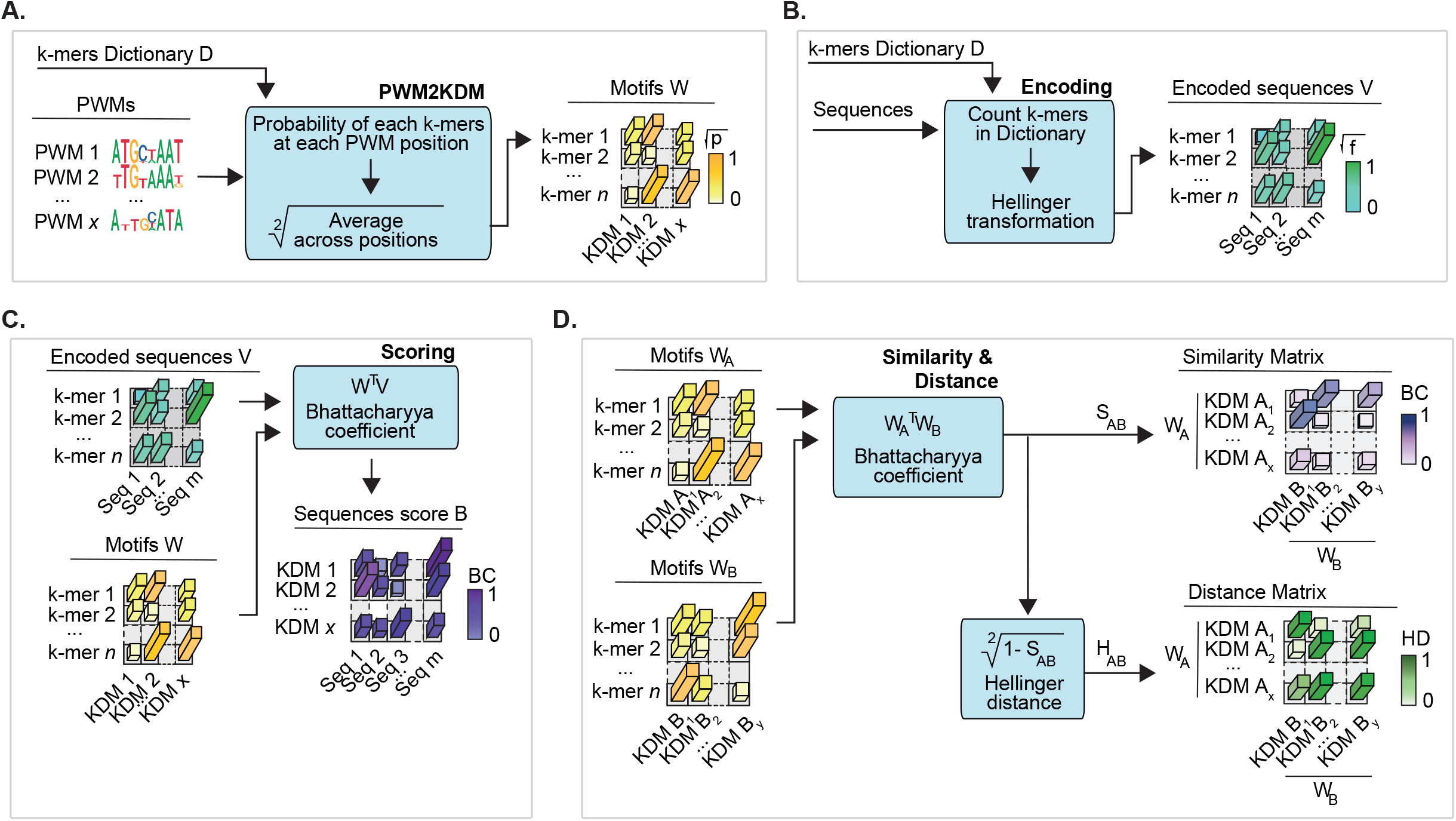
KDM construction, sequence encoding, scoring and motif comparison. **A**. PWM2KDM strategy to derive KDMs W from a set of PWMs using a dictionary D of k-mers. **B**. Sequence encoding as k-mer composition over D using the Hellinger transform. **C**. Scoring KDMs in encoded sequences using the Bhattacharyya coefficient. **D**. Comparing two sets of KDMs by motif–motif similarity (BC) and distance (HD).

#### Sequence Encoding

We encode a sequence by its k-mer composition over *D*, applying the same Hellinger transform used for motifs (**Fig. 2B**). For each dictionary entry *i*, we count occurrences in the sequence and convert them to a relative frequency *q*_*i*_, then take the square root to obtain *v*_*i*_ as in (3). An encoded sequence is thus a column vector *v* of length *n*, the observed counterpart of the motif vector *w*, and a set of *n*□ sequences forms a matrix *V*. Because motifs (*w*) and sequences (*v*) now live in the same square-root space, they can be compared directly.

#### KDM score in sequences

Given a set of encoded sequences *V* and a set of KDMs *W*, the score *b* of a motif *w* and a sequence *v* is defined as their BC (**Fig. 2C**):

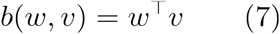

The score is therefore the geometric overlap between the motif’s probable k-mers and the sequence’s observed k-mers. It is high when the k-mers expected under the motif are also frequent in the sequence, and zero when they do not co-occur. For full sets *W* and *V* this generalizes to a single matrix product:

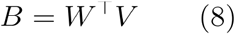

#### Motifs similarity and distance

The same primitive compares motifs to one another. Given two KDM sets *W*_*a*_ and *W*_*b*_, for each pair of motifs *w*_*a*_ and *w*_*b*_ we define their similarity *s*_*ab*_ and distance *h*_*ab*_ as their BC and HD, respectively (**Fig. 2D**). Because both motifs are square-root probability distributions over *D, s*_*ab*_ *= w*_*a*_ ^*T*^*w*_*b*_ measures how much their probable k-mers overlap. For the whole sets we write:

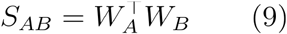

and

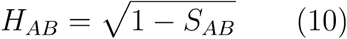

#### KDM Information Content

We measure how specific a KDM *w* is relative to a uniform background over *D* using its IC defined from the Shannon Entropy ^21^ as the maximum uncertainty to which we subtract the KDM uncertainty resulting in the following equation:

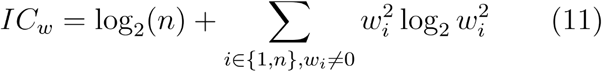

where 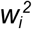 is the probability of k-mer *i*. A motif concentrated on a few k-mers has high IC (low uncertainty), whereas one spread evenly across the dictionary approaches the uniform background and has IC near zero.

#### Merging KDMs or encoded sequences

To merge *m* KDMs (or encoded sequences) we compute, for each k-mer, the square root of the average probability (or frequency):

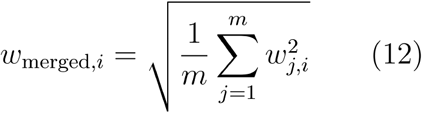

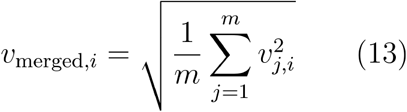

Because the averaging is done on the probabilities 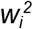 (not on the square-root values *w*_*i*_) before the square root is restored, the merged KDM corresponds to an even mixture of the underlying distributions.

### KDMMap: regional concentration and enrichment of KDMs

When a PWM captures the sequence preferences of a nucleic-acid-binding protein, its strongest matches are typically not uniformly distributed along functionally relevant genomic regions ^22^. Instead, high-scoring sites often cluster at specific positions, such as near ChIP-seq peak summits, cross-link sites, or splice junctions. PWM-based methods exploit this by quantifying the positional enrichment of motif matches. We provide an analogous approach for KDMs.

KDMMap takes as input sets of sequences of equal length aligned to biologically meaningful sites and labeled as positive or negative according to experimental or functional criteria (e.g., bound vs. unbound by a protein, alternative vs. constitutive exons). For each KDM, it tests whether the strongest matches in positives are (*i*) concentrated within a user-defined region and (*ii*) more frequent than in negatives. For each sequence, KDMMap slides a fixed-length window, scores the KDM in each window, and retains the maximum score and its position. It then identifies a score threshold that maximizes classification accuracy between positive and negative sequences. Maximum-scoring positions exceeding this threshold are called hits, yielding at most one hit per sequence, and hit counts at each position are tallied separately for the positive and negative sets (**Fig. 3A**).

**Figure 3.**
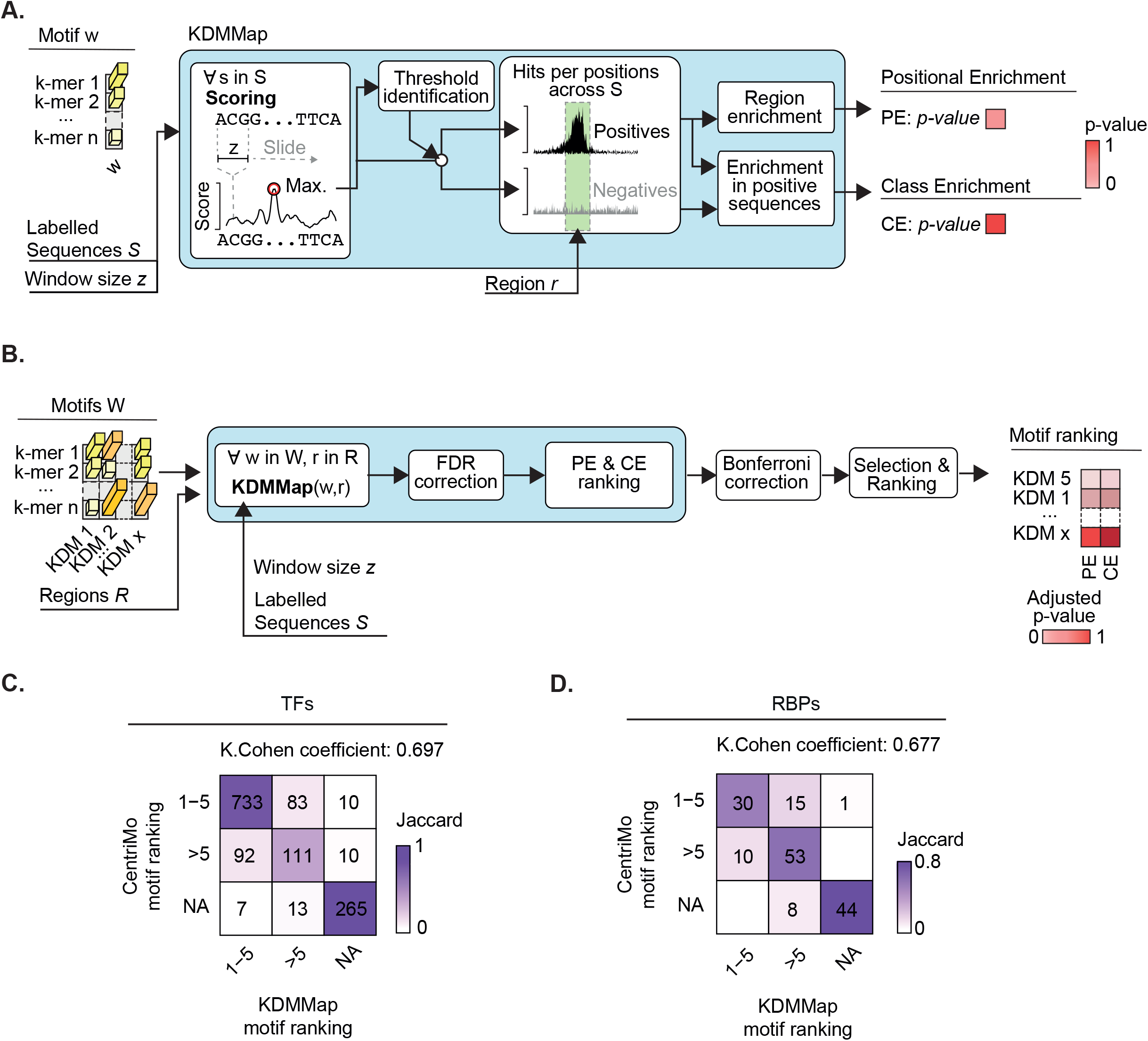
KDMMap workflow and concordance with CentriMo. **A**. KDMMap detects regional concentration (Positional Enrichment, PE) and enrichment (Class Enrichment, CE) of KDMs in sets of labeled sequences. **B**. Workflow for selecting and ranking KDMs based on PE and CE across candidate regions. **C, D**. Pairwise Jaccard similarity matrices comparing CentriMo-ranked PWMs and KDMMap-ranked KDMs for TFs (C, Cohen’s κ = 0.697) and RBPs (D, Cohen’s κ = 0.677). Numbers indicate experiment counts.

To quantify hit concentration in a region of length *L*_*r*_ within sequences of length *L*_*s*_, we use the null expectation of uniform distribution *p*_*0*_ *= L*_*r*_ */L*_*s*_. If positive sequences contain *H*_*r,pos*_ hits inside the region out of *H*_*s,pos*_ total hits, we apply an upper-tail binomial test (*p = p*_*0*_) and summarise the effect size as the Positional Enrichment (*PE*):

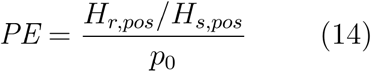

To quantify enrichment of regional hits in positives relative to negatives, we set *p*_*0*_ = *H*_*r,neg*_*/N*_*neg*_ where *H*_*r,neg*_ is the number of negative sequences with a hit in the region and *N*_*neg*_ is the number of negative sequences. We apply an upper-tail binomial test with successes *H*_*r,pos*_ and trials *N*_*pos*_, and define Class Enrichment (*CE*) as:

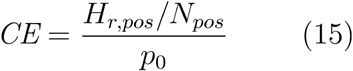

For both *PE* and *CE* we report the corresponding p-values. KDMMap takes as input labeled sequences, a KDM, the region of interest, and the scoring window length (**Fig. 3B**).

We benchmarked KDMMap in a setting where PWM-based tools are widely used. In ChIP- and CLIP-seq, peak calling identifies genomic intervals enriched for sequenced tags. However, these intervals are often broader than the underlying binding site and may contain motifs for co-binding or competing factors ^23,24^. Motifs recovered by *de novo* discovery are therefore not necessarily those directly recognized by the target protein. CentriMo addresses this by ranking PWMs whose predicted sites are centrally enriched within peaks, under the expectation that true target sites cluster around the summit ^22^.

To benchmark KDMMap against CentriMo, we focused on the *PE* task also requiring a significant *CE* (see Methods). We ran CentriMo on 1,324 TF ChIP-seq experiments for which a cognate PWM was present in the HOCOMOCO v13 collection (1,611 PWMs) ^25^. Each experiment was classified by whether the target PWM ranked in the top five, outside the top five, or was not reported. We then converted the HOCOMOCO PWMs into KDMs using PWM2KDM and ran KDMMap on the same experiments, testing all KDMs for *PE* and *CE* across sub-sequences centered on the peak summit. Multiple testing was controlled using Benjamini-Hochberg correction, with an additional Bonferroni correction across KDMs. KDMs were ranked by the most significant region’s adjusted p-values for *PE* and *CE*, and experiments were classified using the same criteria as CentriMo. The two methods agreed strongly, with 84% of experiments (n = 1,109) concordantly classified (**Fig. 3C**). We obtained comparable results for RNA-binding proteins using 161 eCLIP experiments from the ENCODE data portal ^26^, with 1,071 corresponding PWMs from the mCrossBase database ^27^. Agreement between CentriMo and KDMMap remained high, with 79% of experiments (n = 127) concordantly classified (**Fig. 3D**).

Overall, KDMMap provides a KDM-based analogue of established PWM-centric enrichment analyses, enabling quantitative tests of positional clustering and differential enrichment in aligned sequence sets. Across TF ChIP-seq and RBP eCLIP-seq benchmarks, KDMMap closely matched CentriMo’s results, supporting its use for prioritizing candidate motifs that likely reflect direct binding of the assayed protein.

### KDMMatch: information content-aware KDM matching and annotation

Motifs encoding the binding preferences of related proteins often resemble each other, reflecting similarities in their DNA- or RNA-binding domains. This enables functional annotation of an uncharacterized motif (query) by comparison to an annotated reference collection (targets). However, query-target similarity ranks alone are difficult to interpret without a target-specific null model, because some motifs are intrinsically generic and tend to appear similar to many others.

To build an appropriate null distribution for each reference KDM, we computed its similarity to all other motifs in the reference set using the BC defined above. These similarities depended strongly on the IC of the motifs. After converting HOCOMOCO^25^ PWMs to KDMs, the average similarity of a motif to the rest of the collection decreased almost linearly with IC (Pearson correlation coefficient = -0.98, p-value<10^−16^, **Supplementary Fig. S2**). In practical terms, low-information motifs on average show higher similarity with the rest of the collection, inflating their apparent similarity.

Based on these observations, we developed KDMMatch. For each reference motif, we estimate the BC similarities with all other KDMs as a function of IC by fitting a linear trend (**Fig. 4A**, upper part). We convert observed similarities into residuals (the difference between observed and fitted), which are centered at zero by construction. We approximate the residual distribution for each reference motif with a normal distribution *N(0, σ)*, where *σ* is estimated from the residuals of that motif against all other reference KDMs. This provides a KDM-specific null distribution. Given a query KDM *w*_*q*_, we compute its similarity to every reference KDM, subtract the IC-based expectation to obtain a residual for each target, and assign an upper-tail p-value from the corresponding null distribution (**Fig. 4A**, lower part). We correct for multiple testing across targets using Benjamini-Hochberg to obtain q-values and report significant matches ordered by decreasing similarity.

**Figure 4.**
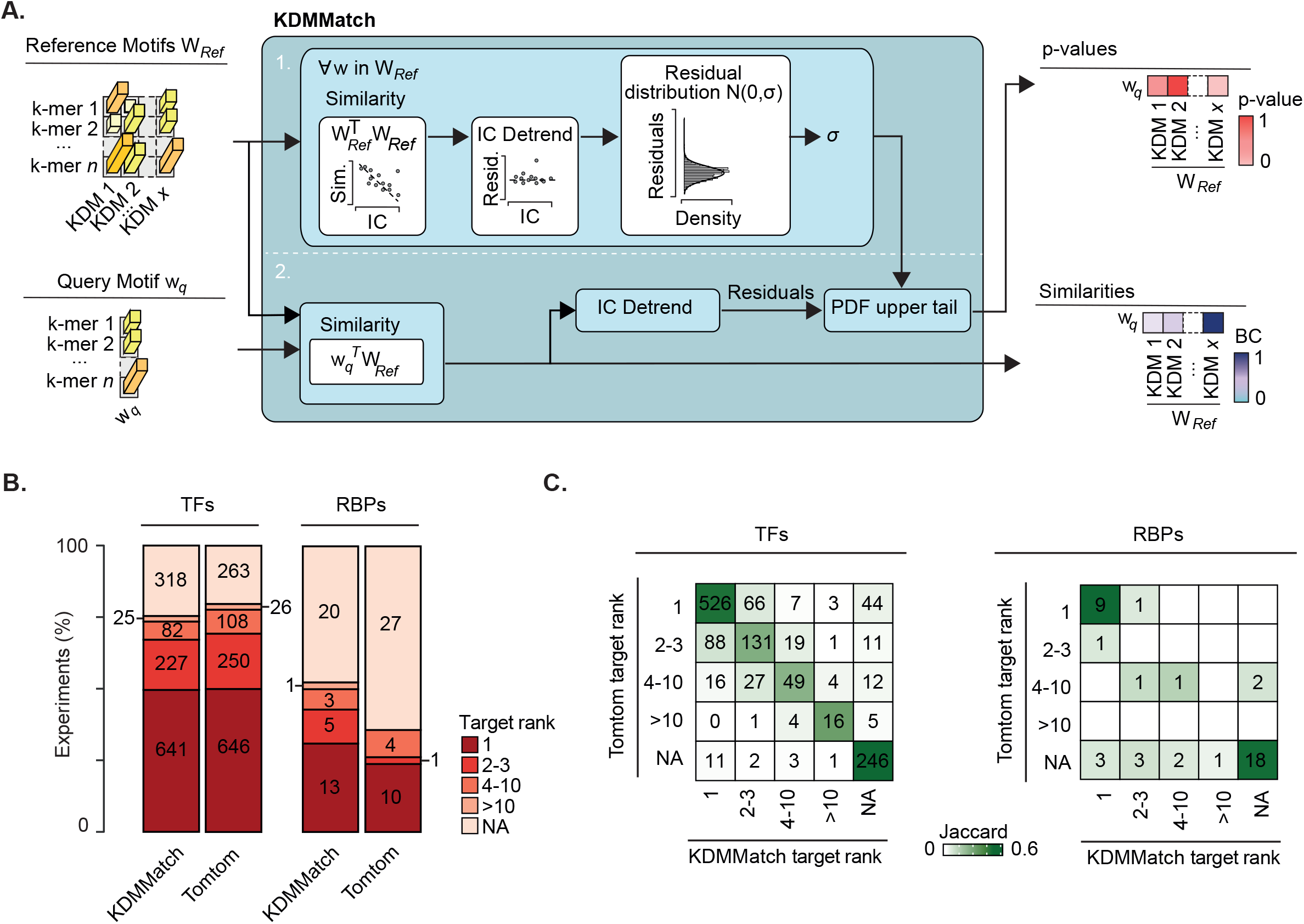
IC-adjusted motif matching with KDMMatch and benchmarking against Tomtom. **A**. KDMMatch reports Bhattacharyya similarity between a query KDM and a reference KDM set, with significance estimates from information-content (IC) adjustment via residuals and KDM-specific null distributions. Step 1: build per-target null. Step 2: score query and assign p-values. **B**. Fraction of TF ChIP-seq (left) and RBP CLIP-seq (right) experiments in which the expected query–target association is ranked as top hit (1), top 3 (2-3), top 10 (4-10), below top 10 (>10), or not found (NA), for KDMMatch and Tomtom. **C**. Pairwise Jaccard similarity matrices comparing Tomtom and KDMMatch query–target rankings for TFs and RBPs. Numbers indicate experiment counts.

We benchmarked KDMMatch against Tomtom from the MEME Suite ^28^, a widely used PWM-based motif-matching tool. For each TF and RBP, we treated experimentally derived motifs as queries and asked whether top-ranked matches recovered the reference motif annotated for that protein. Because both methods produce ranked outputs, we summarized performance by the best rank at which the correct association was recovered, grouping results into five mutually exclusive rank bins (1, 2-3, 4-10, >10, not found).

For TFs, we used query motifs from 1,932 experiments in the Factorbook database ^29^ and the HOCOMOCO collection ^25^ as the reference set. For RBPs, we used motifs from multiple experiments for 42 proteins in CisBP-RNA and ATtRACT databases ^30,31^ with mCrossBase ^27^ as reference.

Both tools detected 50% of expected query-target associations for TFs as top hits, and ∼75% within the top 10 matches (**Fig. 4B**). For RBPs, KDMMatch and Tomtom recovered 31% and 24% of correct associations as top-ranked matches, respectively. Proteins recognize signals with distinct compositions in RNA and DNA. PWMs derived from DNA binding are generally longer and more informative, whereas those associated with RNA binding tend to be shorter and more degenerate (median IC: 0.948 for DNA, 0.861 for RNA, normalized per position). A contingency analysis showed that both approaches assigned most experiments to the same rank category for both TFs (968/1,293; 74.5%) and RBPs (28/42; 66.7%) (**Fig. 4C**), indicating that KDMMatch provides comparable annotation performance while operating directly on the KDM representation.

Together, these results confirm that accounting for information content is critical for interpreting motif similarity and assigning statistically meaningful matches. KDMMatch provides an IC-aware, distribution-based matching strategy that enables robust annotation of KDM motifs with performance comparable to established PWM-based tools.

### KDMFind: *de novo* unsupervised KDM discovery

Consider a set of encoded sequences *V* and a set of *x* known motifs whose patterns are described as a set of *x* KDMs *W*, each present in a sufficiently high number of sequences. From the definitions above we compute a score matrix *B = W*^*T*^*V*, where each entry indicates how similar a KDM’s k-mer probability distribution is to the k-mer frequencies observed in a sequence. The contribution of a given KDM to a square-rooted k-mer frequency in a sequence can be reasonably considered as proportional to the square-rooted probability of that k-mer in the KDM, weighted by the KDM’s score in that sequence. The k-mer *i* square-rooted frequency observed in sequence *j* is then approximated by summing contributions over all KDMs:

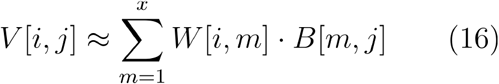

For all sequences we thus can approximate their encoding matrix *V* as *V* ≈ *WB = WW*^*T*^*V*. If *W* is unknown we find it by minimizing the Frobenius norm of the residual:

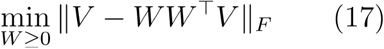

subject to *W* ≥ *0* elementwise and each of its x columns (the KDMs) having unit Euclidean norm 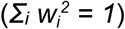, so that each column formally represents a KDM.

This optimization, introduced by Yang et al. ^32^ as PNMF, is solved by a recursive algorithm that yields a non-negative matrix *W* with high sparsity, locality, and orthogonality. The algorithm requires an initial matrix *W*_0_, which is iteratively updated until convergence. Because the problem is non-convex, local minima can be reached depending on *W*_0_. To overcome this, we compute *W*_0_ by applying hierarchical clustering to the dendrogram obtained from *V* by calculating Hellinger distances between its columns. The number of clusters (the factorization rank) can be provided by the user or chosen as the value yielding the highest average silhouette score. The *W*_0_ columns are obtained by merging, for each cluster, the corresponding columns of *V* into a single vector. After convergence, the orthogonality of the solution implies that the columns of *W* tend to be independent, representing distinct motifs. Although these columns have nearly unit norm at convergence, we normalize them as a final step so the solution is formally a set of KDMs (**Fig. 5A**). We named the algorithm KDMFind and applied it to multiple datasets of experimentally identified binding sites for TFs and RBPs (see Methods).

**Figure 5.**
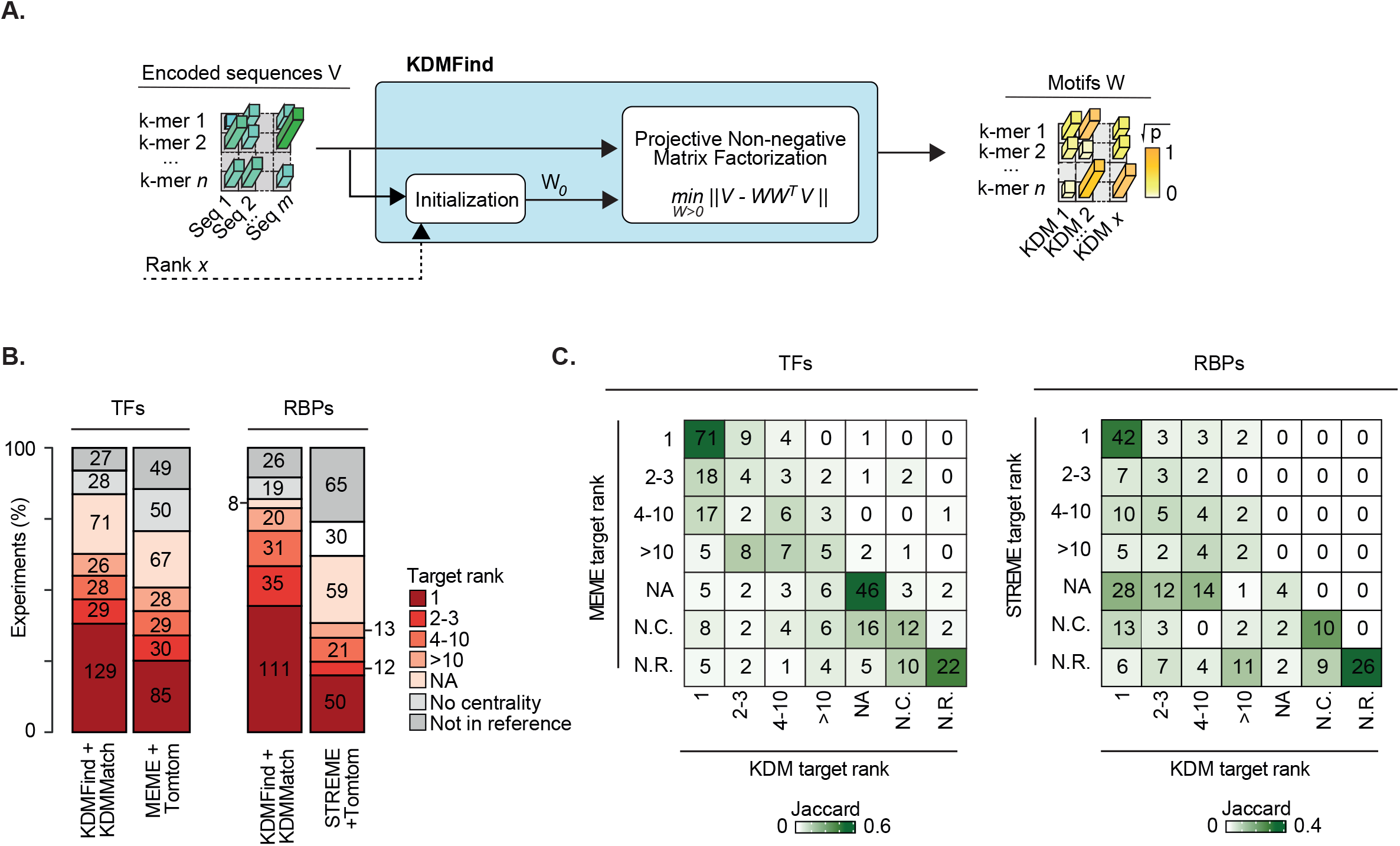
KDMFind workflow and benchmarking against MEME and STREME. **A**. KDMFind applies Projective Non-negative Matrix Factorization (PNMF) to identify de novo KDMs from a set of sequences encoded under a chosen k-mer dictionary. **B**. Counts of experiments by best target rank for novel motifs identified by KDMFind+KDMMatch vs. MEME+Tomtom (TFs, left) or STREME+Tomtom (RBPs, right). Ranks: 1, 2-3, 4-10, >10, NA, No centrality (N.C.), Not in reference (N.R.). **C**. Pairwise Jaccard similarity matrices comparing target ranks between MEME+Tomtom and KDMFind+KDMMatch (TFs) and STREME+Tomtom and KDMFind+KDMMatch (RBPs). Numbers indicate experiment counts.

*De novo* motif-finding algorithms are commonly applied to such experiments to build reference collections of annotated motifs, against which motifs from other sequences are later classified. Within a given experiment, only motifs with high PE and CE are typically taken to be recognized by the target protein and added to the reference collection ^25,29,33^. For both PWMs and KDMs, we therefore built centrally enriched motif collections and asked how well motifs from independent experiments were classified against them according to their target protein.

For each dataset (TFs and RBPs), we defined two independent subsets: a reference subset used to build the collection and a classification subset used to classify motifs against it. For TFs, we used TF-ENCODE (2,037 experiments distributed non-uniformly across 844 target proteins and 70 cell lines). Because 338 TFs had more than one experiment, we formed the classification subset by selecting one experiment at random per TF, leaving the remaining 1,649 experiments as the reference subset. For RBPs, we used RBP-ENCODE (250 eCLIP experiments covering 168 distinct RBPs in HepG2 and K562). Here only 78 of the 168 RBPs had more than one experiment, and redundant experiments came mostly from different cell lines; we therefore used the training set as the reference subset and the test set as the classification subset.

For both TFs and RBPs (Methods), we first applied KDMFind to every experiment in both subsets, then used KDMMap to select up to five of the most centrally enriched KDMs per experiment ^29^. KDMs selected from reference-subset experiments were pooled into a reference motif collection, against which KDMs from the classification subset were matched with KDMMatch. We followed an analogous procedure for PWMs, using MEME ^2^ (TFs) or STREME ^1^ (RBPs) for motif finding, CentriMo ^22^ for centrality, and Tomtom ^28^ for matching. For each experiment in the classification subset, we obtained a ranked list of matches. Each experiment was labeled independently for PWMs and KDMs as “Not in reference” if its target protein was absent from the reference, “No centrality” if no centrally enriched motif was found, or “Not found” (NA) if no match was found between experiment motifs and a reference motif for the target. Otherwise, experiments were assigned to one of four non-overlapping rank classes (1 = best hit, 2-3, 4-10, >10).

For TFs, KDMFind+KDMMatch recovered the cognate target as the top-ranked match in 129 experiments, compared with 85 for MEME+Tomtom, a 51.8% increase in top-rank recovery (**Fig. 5B**, left). The two pipelines agreed on the rank category for most experiments (**Fig. 5C**, left). For RBPs, KDMFind+KDMMatch recovered the corresponding target as the top hit in 111 experiments versus 50 for STREME+Tomtom (**Fig. 5B**, right), reflecting the greater advantage of the distributional representation when motifs are short and degenerate. Concordance with STREME+Tomtom rank categories was high (**Fig. 5C**, right), with most experiments falling on the diagonal or one bin off. These results indicate that KDMFind, combined with KDMMap-based selection and KDMMatch-based annotation, identifies cognate motifs at least as effectively as established PWM workflows, with a particular advantage for RBPs.

### KDM-LRLM: Binding sites prediction based on KDMs derived features

To distinguish two classes of sequences (e.g., bound vs. unbound) using KDM scores, we introduce KDM-LRLM (KDM-based Lasso-Regularized Logistic Model). Given a set of binary-classified sequences, KDM-LRLM first applies KDMFind to identify a set of KDMs, then defines a set of features per sequence based on these motifs. A logistic-regression model (binomial GLM) with L1 (Lasso) regularization is trained on the feature matrix to distinguish the two classes, simultaneously selecting informative features and identifying optimal regression coefficients (**Fig. 6A**). Because different parts of a sequence can show different probabilities of containing motifs high-scoring positions, KDM-LRLM supports multiple features defined by splitting each sequence into multiple sub-sequences on which motif scores are evaluated (**Fig. 6B**).

**Figure 6.**
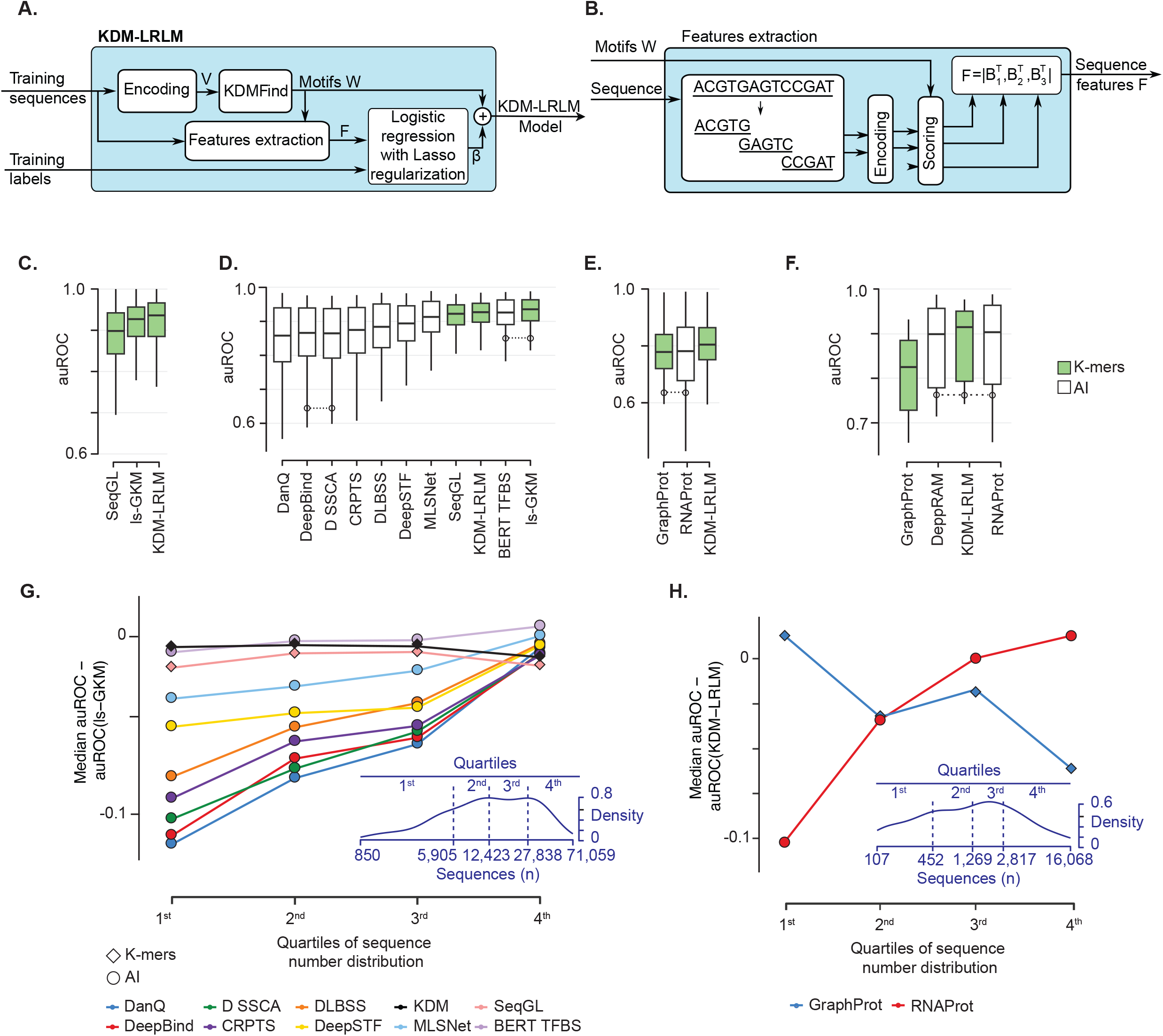
KDM-LRLM workflow and performance benchmarking. **A**. KDM-LRLM workflow: sequences and binary labels fit a binomial GLM with Lasso regularization. **B**. Schematic representation of Features extraction for input to KDM-LRLM. Features are calculated as the scores of a set W of KDMs possibly splitting the sequences in two or more parts. **C**. auROC distributions on TF-ENCODE comparing KDM-LRLM with two k-mer–based methods (green) and AI methods (white). Methods ordered by average rank **D-F**. auROC distributions comparing KDM-LRLM with competing methods across three datasets: TF-benchmark (**D**, ten methods), RBP-ENCODE (**E**, two methods), and RBP-benchmark (**F**, three methods). In all panels, methods are ordered by average rank. Boxplots connected by dashed lines indicate distributions not significantly different (paired Wilcoxon, p > 0.05). **G**. Difference between median auROC of each method and ls-GKM on TF-benchmark, with experiments grouped by training-set size quartiles. Diamonds: k-mer–based; circles: AI-driven. Density plot (bottom right) shows the distribution of sequence counts across experiments; dashed lines mark quartile thresholds. **H**. Difference between median auROC of GraphProt and RNAProt and KDM-LRLM on RBP-ENCODE, grouped by training-set size quartiles. Diamonds: k-mer–based; circles: AI-driven.

We applied KDM-LRLM to four datasets: TF-ENCODE (2,037 experiments), TF-benchmark (165 experiments) ^9,34^, RBP-ENCODE (250 experiments), and RBP-benchmark (23 experiments) ^16^ (see Methods). Predictive performance was evaluated for each experiment using the area under the Receiver Operating Characteristic curve (auROC), the area under the Precision–Recall curve (auPRC), and accuracy (**Supplementary tables S2-S5**). In the main text we focus on auROC (**Fig 6 and Supplementary Fig. S3**); auPRC and Accuracy results, which confirm the auROC trends, are reported in **Supplementary Figures S4-S6**. For RBP-benchmark only auROC is reported, as the original studies did not provide the data required to compute additional metrics.

On TF-ENCODE (301 bp sequences), KDM-LRLM features were defined by splitting each sequence into three overlapping parts (Methods). We compared KDM-LRLM with two k-mer–based methods, ls-GKM^5^ and SeqGL^6^. KDM-LRLM performed significantly better than both (paired one-sided Wilcoxon tests, **Fig. 6C, Supplementary Fig. S3A**).

On TF-benchmark (101 bp), sequences were too short to allow for positional splitting; we therefore defined a single feature per motif in W. In addition to ls-GKM ^5^ and SeqGL ^6^, we obtained performance measures for eight deep-learning tools (BERT-TFBS ^15^, DanQ ^8^, DeepBind ^35^, DeepSTF ^12^, MLSNet ^13^, DLBSS ^11^, D-SSCA ^9^, CRPTS ^10^; see Methods). ls-GKM was the best-performing method overall, followed by BERT-TFBS (not significantly different from ls-GKM), KDM-LRLM, and SeqGL; the remaining AI-based methods performed significantly worse than KDM-LRLM (**Fig. 6D, Supplementary Fig. S3B**).

On RBP-ENCODE (71 bp), we compared KDM-LRLM with a k-mer–based method (GraphProt ^7^) and a deep-learning method (RNAProt ^16^). KDM-LRLM significantly outperformed both, which were statistically indistinguishable from each other (**Fig. 6E, Supplementary Fig. S3C**).

On RBP-benchmark (81 bp), we additionally included DeepRAM^17^ in the comparison. KDM-LRLM performed on par with both RNAProt and DeepRAM, with no statistically significant differences among the three methods, while GraphProt performed significantly worse than all others (**Fig. 6F, Supplementary Fig. S3D**).

We next examined how predictive performance depended on the number of sequences available for training (**Fig. 6G-H**). Across both the TF and RBP datasets, the gap between deep-learning and k-mer-based methods narrowed steadily as training-set size increased, but the two categories reached parity from opposite directions. AI methods improved sharply with more data, whereas k-mer methods were already strong in the small-data size and gained comparatively little.

On TF-benchmark, performance was expressed relative to ls-GKM (**Fig. 6G**). In the smallest-data quartile, the AI methods trailed ls-GKM by a wide margin, with most clustered well below it, while the k-mer methods (KDM-LRLM and SeqGL) tracked ls-GKM near the top. As training-set size grew, the AI methods rose steeply and converged toward the k-mer line, surpassing it only in the largest quartile, where DeepSTF and MLSNet exceeded both SeqGL and KDM-LRLM. BERT-TFBS was the consistent exception. Built on a pre-trained encoder, it performed near the k-mer methods across all quartiles, showing little dependence on the size of the task-specific training set.

The RBP comparison showed the same dependence as a direct crossover (**Fig. 6H**). Relative to KDM-LRLM, the deep-learning method RNAProt rose monotonically from well below KDM-LRLM in the smallest quartile to slightly above it in the largest, while the competing k-mer method GraphProt moved in the opposite direction, declining from near parity to well below KDM-LRLM over the same range. The two methods crossed at intermediate training sizes, so that RNAProt overtook KDM-LRLM only where data were most abundant.

Together, these patterns indicate that the recent dominance of deep learning in motif analysis reflects access to large training sets at least as much as architectural innovation. For the majority of real-world binding-prediction experiments, where only a few thousand bound sequences are available, k-mer–based representations such as KDMs remained competitive, and often preferable, and the principal exception, BERT-TFBS, owed its small-data robustness to pre-training rather than to a task-specific architecture.

## Discussion

We introduced KDM, a unifying framework in which motifs and sequences are represented as probability distributions over a shared k-mer dictionary and compared using a single primitive, the Bhattacharyya coefficient. This common geometry enables the four canonical tasks of motif analysis, namely scoring, enrichment, annotation, and discovery, to be performed within one consistent representation, and integrates naturally with downstream binding prediction via KDM-LRLM. Across more than 2,000 TF ChIP-seq and 300 RBP CLIP-seq experiments, KDM-based tools match or exceed the performance of established PWM-based and machine-learning alternatives while retaining interpretability.

A defining feature of KDM-LRLM is that its predictive features are derived from motifs that are themselves discoverable, scorable, and classifiable within the same framework, a property none of the methods we compared to share. In deep-learning predictors, feature extraction and classification are entangled in a single model, and interpretable motifs are reconstructed from learned parameters as approximate PWMs. K-mer SVMs are transparent but classify on individual k-mer counts, which are not motifs so cannot be annotated. In KDM-LRLM, by contrast, the features entering the Lasso model are directly obtained from KDMs that can be enriched (KDMMap), annotated (KDMMatch), and compared to reference collections directly, so a prediction can be traced back to specific, named motifs. This matters because PWM-based predictors, despite their interpretability, perform poorly as classifiers relative to k-mer and deep-learning models ^36^. KDM-LRLM occupies the otherwise sparse middle ground, retaining motif-level interpretability while remaining competitive with the best non-PWM predictors.

A key conceptual feature of the KDM representation is that, unlike a PWM, it does not commit to a single positional alignment. A KDM can capture both classical localized motifs (where one or a few k-mers dominate the distribution) and “diffuse” multivalent binding preferences, in which recognition arises from the co-occurrence of several short, possibly degenerate elements within a region ^3,37^. For RNA-binding proteins, where such multivalent signatures are common, this property likely explains the substantial advantage of KDMFind+KDMMatch over STREME+Tomtom ^1,28^in recovering cognate motifs (111 vs. 50 top-rank matches, **Fig. 5B**). For TFs, the advantage is more modest but consistent (129 vs. 85), suggesting that even canonically localized binding can benefit from a distributional treatment when motif length or degeneracy varies across factors.

A second, more surprising finding concerns the relative performance of deep-learning and k-mer-based binding predictors. In the TF-benchmark, only one AI method built around a pre-trained encoder (BERT-TFBS ^15^) is competitive with the best k-mer methods overall; the remaining seven AI methods are matched or exceeded by KDM-LRLM, SeqGL ^6^, and ls-GKM ^5^. Stratifying by training-set size shows that AI methods overtake k-mer methods only in the top quartile of experiments. Notably, BERT-TFBS was the single AI method to remain competitive in the small-data regime, and it is also the only one built on an encoder pre-trained on large unlabeled sequence datasets, precisely the feature that reduces dependence on task-specific training data. This is suggestive, because where the other deep models must learn their representation from each experiment’s labeled sequences, BERT-TFBS inherits a general sequence representation and need only fit a lightweight task head. The same logic applies, in a different form, to KDM-LRLM. KDMFind performs an unsupervised, label-free factorization of the sequence set before any classifier is trained, so the KDM features are effectively a self-supervised “pre-training” step that supplies a strong representation in the low-data size. Viewed this way, the two methods that hold up when data are scarce, BERT-TFBS and KDM-LRLM, are the two that decouple representation learning from the supervised objective.

Beyond binding prediction, KDMFind offers a route to identify motifs in large sequence sets even when they occur in narrow regions, because PNMF identifies recurrent k-mer compositions independent of where they appear within sequences. This property is potentially useful for applications such as splice-site analysis, enhancer dissection, and the discovery of cis-regulatory grammars in long sequences, which we plan to explore in future work.

Several limitations should be noted. First, the size of the k-mer dictionary grows rapidly with *k*, and double-stranded DNA dictionaries with two gaps already comprise 57,472 entries at *k* = 8; larger *k* or more permissive gap structures would require sparse representations and parallel computation, which our implementation already exploits. Second, KDMFind requires either a user-specified rank or an automatic choice based on silhouette score; for very large datasets we use a fixed rank, which trades adaptivity for tractability. Third, although the IC-aware null in KDMMatch is calibrated empirically per reference motif, it assumes approximate normality of residuals; sensitivity to this assumption merits further study. Finally, the KDM representation is by construction agnostic to positional information within a motif; for proteins whose specificity is dominated by strict positional constraints (e.g., zinc-finger arrays binding long, well-aligned sites), a PWM may remain the more informative representation, and the PWM2KDM conversion is a natural bridge between the two.

Together, these results position KDM not as a replacement for PWMs but as a complementary representation that scales naturally from interpretable motif annotation to competitive binding prediction, within a single mathematical framework.

## Methods

### Citation analysis

To measure how each method is used across research fields, its citations were analysed. For every method in Supplementary Table S1, the original article(s) designated by the authors as the reference to cite when using the method were first identified. All papers citing these articles (published between 2001 and November 2025) were then retrieved from Scopus. To capture the research field of each citing paper, it was labelled according to the subject category of the journal in which it appeared, using the Web of Science (WoS) journal categories downloaded in November 2025. The seven categories represented across all methods were retained: “Computer Science, Artificial Intelligence”, “Computer Science, Interdisciplinary Applications”, “Mathematical & Computational Biology”, “Biochemical Research Methods”, “Biotechnology & Applied Microbiology”, “Biochemistry & Molecular Biology”, and “Genetics & Heredity”. Finally, for each method, the number of distinct papers citing it within these categories was counted, along with the fraction falling into each.

### Datasets and preprocessing

Multiple publicly available ChIP-seq, CLIP-seq, and eCLIP-seq datasets were assembled to evaluate KDM-based motif analyses across DNA- and RNA-binding proteins. Datasets were selected to cover three complementary benchmarking scenarios: large-scale ENCODE collections used in prior motif discovery resources ^26^, established TF binding site prediction benchmarks, and widely used RBP cross-linking benchmarks. For each dataset, positive sequences were defined from experimentally supported binding regions or cross-link sites; matched negative sequences were generated as described below. All datasets were split into training and test sets as specified for each analysis.

#### TF-ENCODE ChIP-seq dataset

A total of 2,037 ChIP-seq experiments matching those used by Factorbook ^29^ to derive PWMs were retrieved from the ENCODE data portal ^26^, restricting to experiments available as of July 14, 2022 and mapped to GRCh38. This freeze was chosen to match the Factorbook PWM-derivation cohort ^29^ and ensure comparability between PWM- and KDM-based analyses. For each experiment, peaks were ranked by signal value and up to the top 1,000 were retained; a 301 bp sequence centered on each peak summit was extracted to form the positive set. Matched negative sequences were generated by randomly sampling genomic regions that did not overlap any peak from the same experiment and matched the positive sequences in length, GC content, and repetitive-element content. Positive and negative sequences were then randomly split into training and test subsets.

RBP-ENCODE eCLIP-seq dataset. eCLIP-seq NarrowPeak BED and bigWig files were retrieved from the ENCODE data portal ^26^. IDR-concordant cross-linked site clusters were selected and mapped to GRCh38, yielding 250 experiments covering 168 RBPs across two cell lines (K562 and HepG2). For each cluster, the summit position was defined as the average of the replicate signal maxima within the cluster, computed from replicate-specific bigWig profiles. Positive sequences were generated by extending 35 bp upstream and downstream of each summit, yielding 71 bp regions. This length was chosen to support partitioning into three 31 bp windows with 11 bp overlap ^3^, thereby accommodating k-mers spanning window boundaries. Negative sequences were generated by randomly sampling genomic regions that did not overlap any positive sequence from the same experiment and matched the positive sequences in length, GC content, and repetitive-element content. For each experiment, sequences were randomly split into training (50%) and test (50%) subsets.

#### TF-benchmark dataset

A total of 165 ChIP-seq experiments spanning 29 TFs across 32 cell lines were obtained from Zeng et al. ^34^. Positive sequences correspond to DNA regions experimentally validated to contain TF binding sites; negative sequences were generated by dinucleotide-preserving shuffling of the corresponding positives. All sequences are 101 bp long. For each experiment, positive and negative sequences were split into training (80%) and test (20%) subsets.

#### RBP-grid and RBP-benchmark datasets

Both datasets were obtained from ^16^. In both cases, sequences are length-normalized to 81 nt, with equal numbers of positive sequences (containing cross-link sites) and negative sequences (without cross-link sites), and split into training (90%) and test (10%) subsets. The RBP-grid dataset comprises 30 eCLIP experiments covering 30 RBPs retrieved from the ENCODE data portal ^26^, with each experiment containing between 6,000 and 10,000 positive sequences. This dataset was used to optimize the k-mer dictionary parameters and factorization rank for RBP analyses by grid search (see below). The RBP-benchmark dataset comprises 23 PAR-CLIP, iCLIP, and HITS-CLIP experiments covering 20 RBPs, originally assembled by Maticzka et al. (2014) for the GraphProt publication ^7^. Each experiment contains up to 5,000 positive sequences. This dataset was used to evaluate the predictive performance of KDM-LRLM against published methods.

### PWM collections for TFs and RBPs

Reference PWM collections were assembled to support PWM-based analyses, PWM-to-KDM conversion, and motif-annotation benchmarks. For TFs, two PWM collections were used. The HOCOMOCO v13 in vivo collection ^25^ comprises 1,611 non-redundant PWMs covering 1,120 TFs (lengths 7-31 bp). The Factorbook PWM collection ^29^ was filtered to retain only PWMs derived from the same 2,037 TF-ENCODE ChIP-seq experiments, yielding 6,043 motifs covering 809 TFs (lengths 7-30 bp). For RBPs, three PWM collections were used. mCrossBase ^27^ was filtered to experiments for which at least one PWM was reported, yielding 1,071 PWMs corresponding to 103 RBPs profiled in HepG2 and K562. All mCrossBase PWMs were 11 bp long. CisBP-RNA ^30^ and ATtRACT ^31^ were filtered to Homo sapiens PWMs whose RBPs were also present in mCrossBase, yielding 494 PWMs spanning 42 RBPs (lengths 4-25 bp).

### Definition of k-mer dictionaries and factorization ranks for RBPs

For RBP analyses, *L, G*, and the factorization rank *R* used for motif discovery were jointly optimized by grid search on RBP-grid, a dedicated dataset comprising 30 eCLIP experiments. The following combinations were evaluated: *L* ∈ {4, 6}, *G* ∈ {0, 1, 2, 3}, and *R* ∈ {10, 20, 40, 70, 100}. For each combination, positive and negative sequences from each experiment were encoded and KDM-LRLM was applied to the training set; performance was assessed as auROC on the test set. The combination *L* = 6, *G* = 2, *R* = 100 yielded the highest median auROC across experiments and was adopted for all subsequent RBP analyses (**Supplementary Fig. S1, Supplementary Table S6**).

### Benchmarking KDMMap against CentriMo

CentriMo ^22^ and KDMMap were compared on matched motif collections and sequence sets, using harmonized input definitions and consistent multiple-testing correction throughout. For TF analyses, both methods were applied to TF-ENCODE using HOCOMOCO v13 ^25^ as the motif reference. CentriMo was run with default parameters and negative control sequences supplied via --neg. KDMMap was run with halfWin=15, a window size sufficient to accommodate the longest PWM in HOCOMOCO v13. For RBP analyses, both methods were run using mCrossBase PWMs ^27^. KDMMap was run with halfWin=6 to accommodate, whereas, CentriMo was run with --local and --norc and negative control sequences were supplied via --neg; all remaining parameters were left at their default values. The --local flag enables detection of motifs enriched near, but not necessarily at, the sequence center, while --norc disables reverse complement matching, as appropriate for single-stranded RNA sequences.

Because CentriMo reports significance for centrality but not for Class Enrichment (CE), CE p-values were computed separately for each region called significant by centrality (note that CentriMo’s centrality is analogous to the Positional Enrichment, PE, metric in our KDMMAP method). For each such region, an upper-tail binomial test was performed in R (‘stats::binom.test’; ^38^) with the number of successes equal to the number of positive sites in the region, and the null success probability equal to the fraction of negative sites falling within an equally sized region. CE p-values were converted to q-values by multiplying by the number of significantly central regions. Regions were retained if the centrality E-value was < 0.05, the CE q-value was < 0.05, and the CE fold change was > 1; retained regions were ranked by the sum of their ranks on −log_10_(centrality E-value) and −log_10_(CE q-value). Centrality and CE p-values were FDR-corrected and converted to E-values by multiplying by the number of KDMs tested. Regions were retained if both centrality and CE E-values were < 0.05 and CE fold change was > 1, and ranked by the sum of their ranks on −log_10_(E-value_PE_) and −log_10_(E-value_CE_).

### Motif matching with Tomtom and KDMMatch

For TF analyses, Tomtom ^28^ was run with –thresh 0.05; all remaining parameters were left at their default values. KDMMatch was run with default settings and a q-value threshold of 0.05.

For RBP analyses, Tomtom was run with --xalph, --norc, and --thresh 0.05; all remaining parameters were left at their default values. KDMMatch was run with default settings; matches with q-value < 0.05 were considered significant.

### De novo motif discovery

For TF analyses, MEME ^2^, was run on TF-ENCODE using the following parameters: --revcomp --evt 0.01 --dna. KDMFind was applied to each experiment independently; the factorization rank was automatically selected by maximizing the average silhouette width (ASW) computed on the Hellinger-distance matrix between all encoded sequences, using hierarchical clustering. The resulting sequence clusters were aggregated by averaging the encoded sequences within each cluster to derive the initialization matrix *W*_*0*_.

For RBP analyses, STREME ^1^ was run on RBP-ENCODE with parameters --rna --maxw 12 --nmotifs K, where K = min(100, N) and N is the number of input sequences. KDMFind was applied with the factorization rank determined by the grid search procedure described above.

### Binding sites predictions

Four datasets were used to compare binding site prediction performance across multiple methods. In all cases, models were fitted on a training set and evaluated on a test set; performance was assessed using auROC, auPRC, and accuracy.

We used the TF-ENCODE to compare KDM-LRLM, SeqGL ^6^, and ls-GKM ^5^. For KDM-LRLM, the KDMs obtained through KDMFind during de novo motif discovery (see above) were reused directly. Input features were defined by partitioning each sequence into three 121 bp windows with 31 bp overlap. SeqGL and ls-GKM were fitted with default parameters.

The TF-benchmark dataset was used to evaluate KDM-LRLM against eight AI-based methods whose performance estimates were retrieved from ^13,15^: BERT-TFBS ^15^, DanQ ^8^, DeepBind ^35^, DeepSTF ^12^, MLSNet ^13^, DLBSS ^11^, D-SSCA ^9^, and CRPTS ^10^. Since sequences in this dataset are 101 bp long, a single-window feature approach was used. Because computing the full Hellinger distance matrix for clustering-based initialization scales as O(n^2^), experiment-size-dependent settings were adopted for the factorization rank: for experiments with up to 10,000 sequences, the same procedure as for TF-ENCODE was applied; for experiments with 10,000–20,000 sequences, the rank was fixed to 5,000; for experiments exceeding 20,000 sequences, the rank was fixed to 10,000 and W_0_ was constructed from 20,000 randomly sampled sequences. SeqGL and ls-GKM were fitted with default parameters.

KDM-LRLM, GraphProt ^7^, and RNAProt ^16^ were compared on the RBP-ENCODE. For KDM-LRLM, the KDMs obtained through KDMFind during de novo motif discovery (see above) were reused directly. Input features were defined by partitioning each sequence into three 31 bp windows with 11 bp overlap. GraphProt and RNAProt were fitted with default parameters. Since RNAProt requires a minimum of 100 input sequences, experiments not meeting this threshold were excluded, resulting in a final set of 213 out of 250 experiments.

The RBP-benchmark dataset was used to evaluate KDM-LRLM against GraphProt, RNAProt, and DeepRAM ^17^. For KDM-LRLM, rank 100 was used as determined by grid search (see above), and input features were defined by partitioning each sequence into three 35 nt windows with 11 nt overlap. GraphProt was fitted with default parameters; performance estimates for RNAProt and DeepRAM were retrieved from ^16^.

### Methods implementation and statistical software

Most of the algorithms presented in this paper are implemented as a C++ library which is an extension for the GeCo Library ^39^ and uses CUDA ^40^ and Armadillo ^41,42^ as backbones for parallel computation, relying on sparse matrices to reduce the memory required to represent KDM-encoded sequences. An R package wrapping the library provides the user-facing API and was used for all analyses in this paper. R ^38^ scripts were used to perform all the analyses.

The function cv.glmnet from the R package glmnet ^43^ with family = “binomial”, alpha = 1, and type.measure = “auc” was used for the identification of the logistic lasso regularized models used by KDM-LRLM. Methods performances in terms of auROC, auPRC and Accuracy were calculated using the ROCR R package ^44^. For KDMMatch the R function lm (default parameters) was used to obtain the linear dependence between motifs IC and similarity. For performance comparison the Wilcoxon signed rank test was applied through the R wilcox.test function ^38^.

## Acknowledgments

U.P. and L.F were supported by the Italian Ministry of Health -”Ricerca Corrente” program. M.C. is supported by AIRC under BRIDGE 2023 ID. 28739 project and the Compagnia di San Paolo Foundation. T.B. is supported by the UniverLecco Association. T.B. is a PhD student within the European School of Molecular Medicine (SEMM).

## Data availability

Data, including bed files as well as experiments metadata for both TF-ENCODE and RBP-ENCODE datasets, are available at https://github.com/cereda-lab/KDM.

## Code availability

The original code has been deposited on GitHub and is publicly available at https://github.com/cereda-lab/KDM. Any additional information required to reanalyze the data reported in this paper is available from the lead contact upon request.

