## Supplemental figures for "KDM: embedding DNA/RNA motifs and sequences in a shared k-mer space for unified discovery, analysis and binding prediction"

**Supplementary Figure** **3**
Figure S1. Grid search for the k-mer dictionary and PNMF rank in RBP analyses. 3
Figure S2. Information content vs. average PWM similarity across HOCOMOCO KDM-converted motifs 4
Figure S3. Median differences in auROC between methods across datasets. 5
Figure S4. auPRC and Accuracy results on the TF ENCODE dataset. 6
Figure S5. auPRC and accuracy on TF-benchmark. 7
Figure S6. auPRC and accuracy on RBP-ENCODE. 8

### Supplementary Figure

#### Figure S1. Grid search for the k-mer dictionary and PNMF rank in RBP analyses.

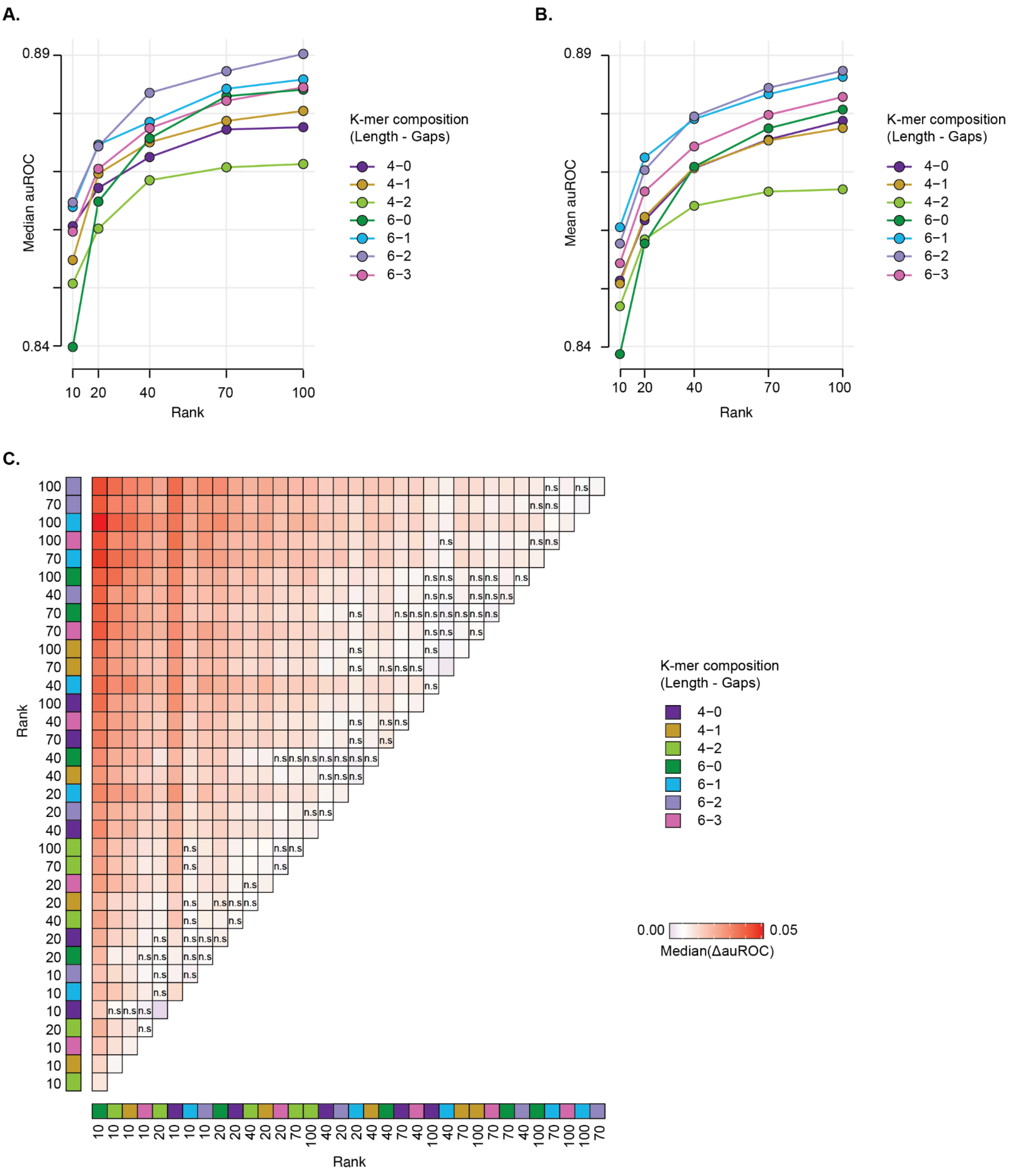


Performance was evaluated on RBP-grid (30 ENCODE eCLIP experiments) under all combinations of k-mer length L ∈ {4, 6}, allowed gaps G ∈ {0, 1, 2, 3}, and PNMF rank R ∈ {10, 20, 40, 70, 100}. For each combination, KDM-LRLM was trained on 90% of the sequences and tested on the remaining 10%. **A**. Median auROC across the 30 experiments for each dictionary × rank combination. **B**. Mean auROC across the 30 experiments for the same combinations. **C**. Matrix of pairwise median auROC differences across combinations; "n.s." denotes pairs not significantly different (paired one-sided Wilcoxon test, $p > 0.05$). p-values are corrected for multiple testing using Bonferroni method. The combination L = 6, G = 2, R = 100 yielded the highest median auROC and was adopted as the default for RBP analyses.

#### Figure S2. Information content vs. average PWM similarity across HOCOMOCO KDM-converted motifs

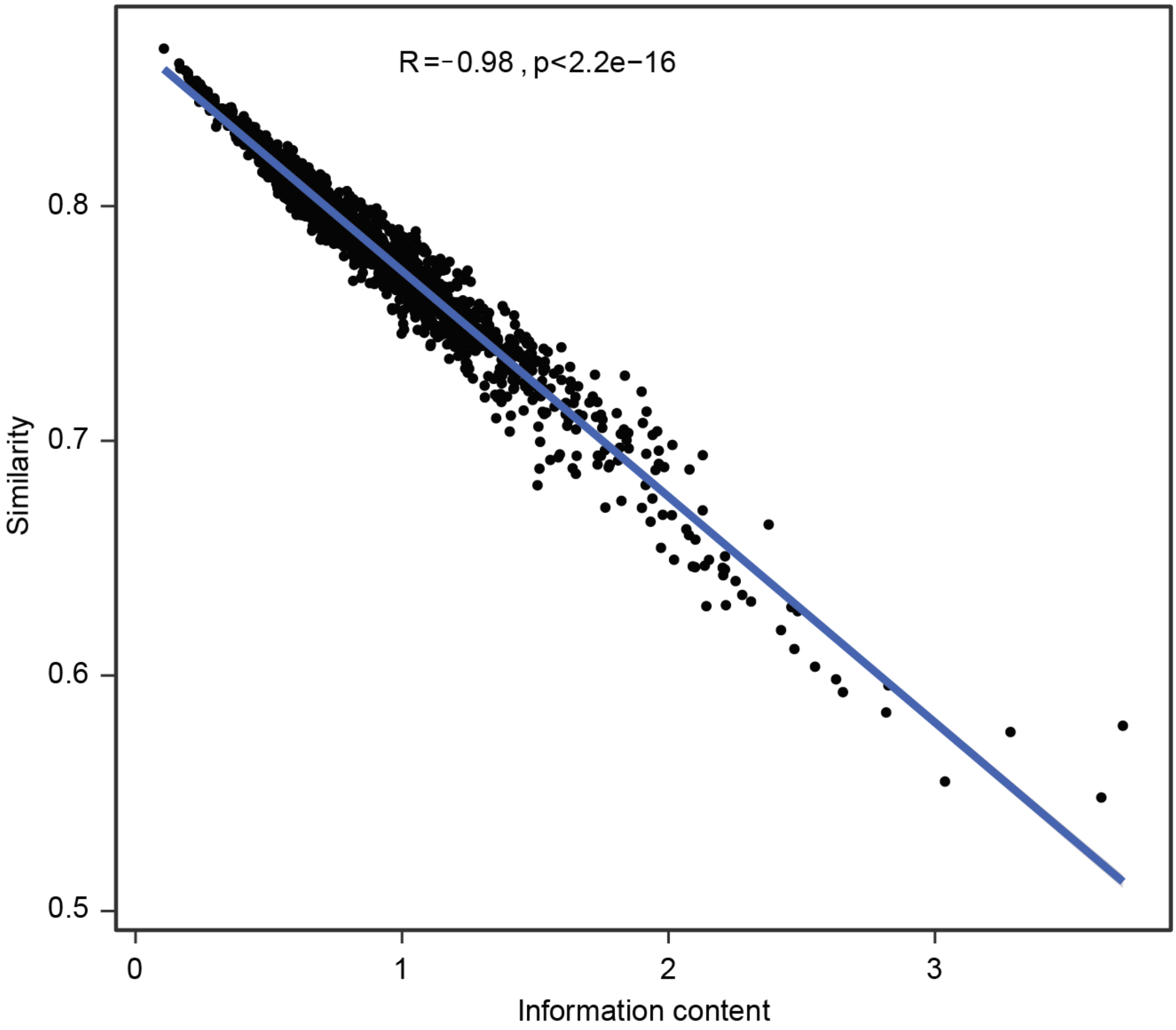


Each dot represents a HOCOMOCO motif converted from PWM to KDM format using PWM2KDM. The blue line represents a linear regression fit ($R = -0.98$, $p < 2.2\times10^{-16}$).

**Figure S3. Median differences in auROC between methods across datasets.**

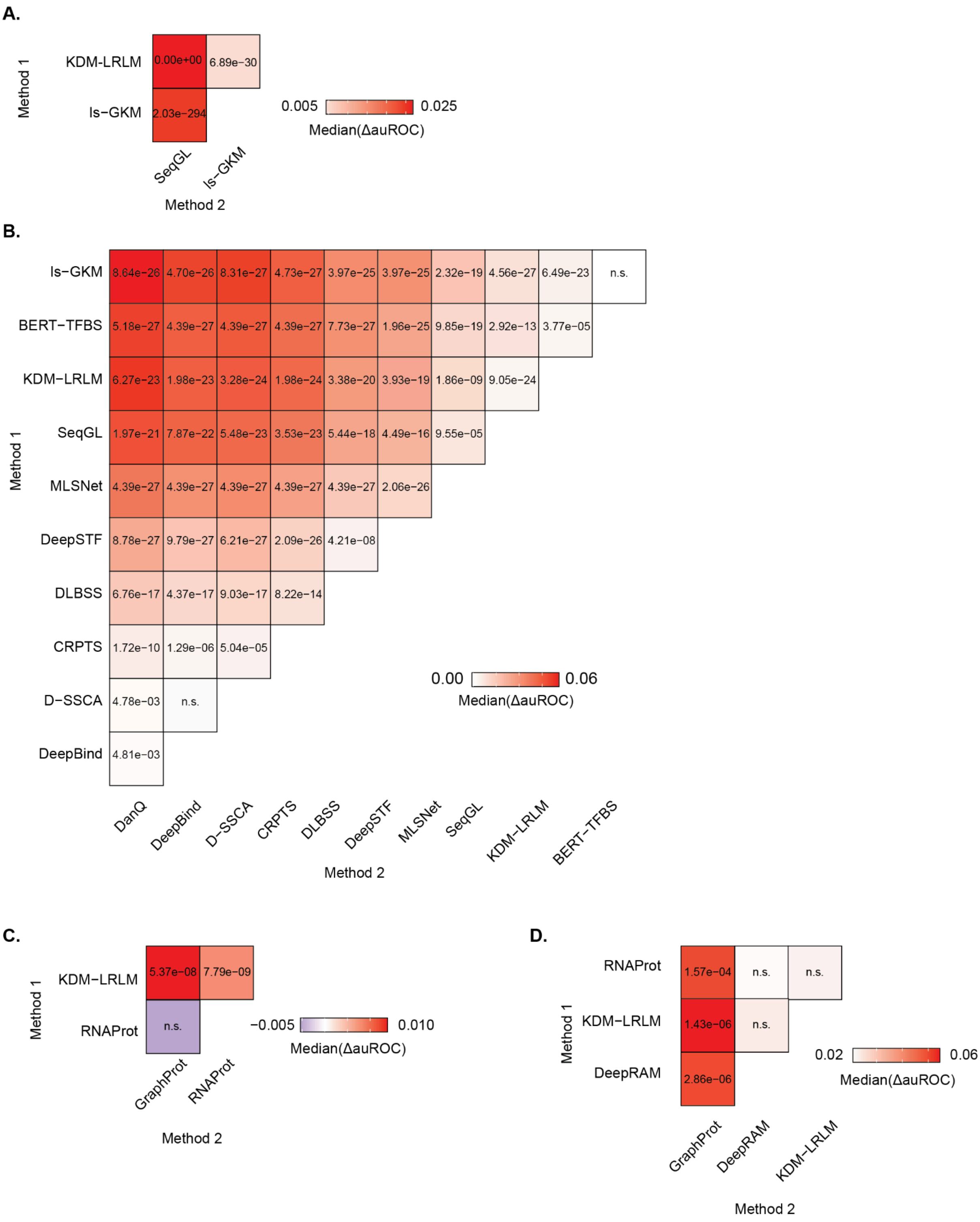


**A.** Matrix of pairwise median auROC differences across the three methods tested on the TF-ENCODE; values indicate p-values corrected for multiple testing (bonferroni) from paired one-sided Wilcoxon tests, "n.s." denotes p > 0.05. **B.** Matrix of pairwise median auROC differences across the eleven methods tested on the TF-BENCHMARK dataset; values indicate p-values corrected for multiple testing (bonferroni) from paired one-sided Wilcoxon tests, "n.s." denotes p > 0.05. **C.** Matrix of pairwise median auROC differences across the three methods tested on the RBP-ENCODE; values indicate p-values corrected for multiple testing (bonferroni) from paired one-sided Wilcoxon tests, "n.s." denotes p > 0.05. **D.** Matrix of pairwise median auROC differences across the four methods tested on the RBP-BENCHMARK dataset; values indicate p-values corrected for multiple testing (bonferroni) from paired one-sided Wilcoxon tests, "n.s." denotes p > 0.05.

#### Figure S4. auPRC and Accuracy results on the TF ENCODE dataset.

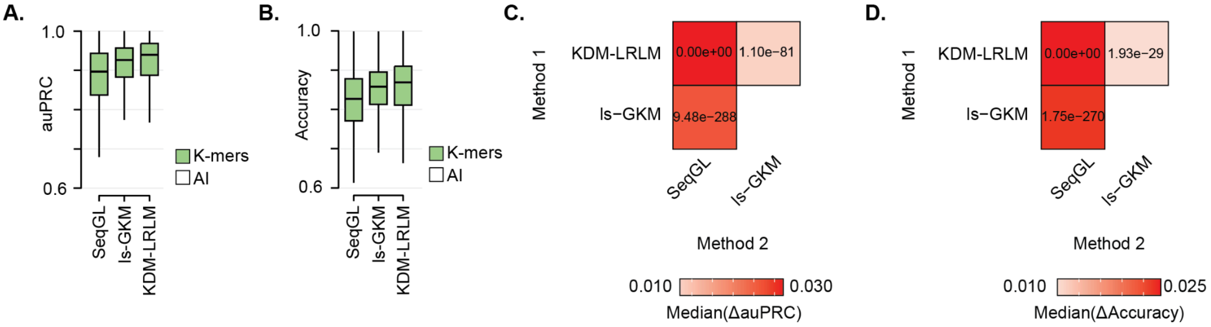


Companion to **Fig. 6C**, with the area under the Precision–Recall curve (auPRC) and Accuracy as evaluation metrics. **A**. auPRC distributions across TF-ENCODE for KDM-LRLM, ls-GKM, and SeqGL. Green: k-mer–based methods; white: AI-driven (none in this panel). **B**. Accuracy distributions for the same comparison. **C**. Matrix of pairwise median auPRC differences across the three methods; values indicate p-values corrected for multiple testing (bonferroni) from paired one-sided Wilcoxon tests, "n.s." denotes p > 0.05. **D**. As in C for Accuracy.

#### Figure S5. auPRC and accuracy on TF-benchmark.

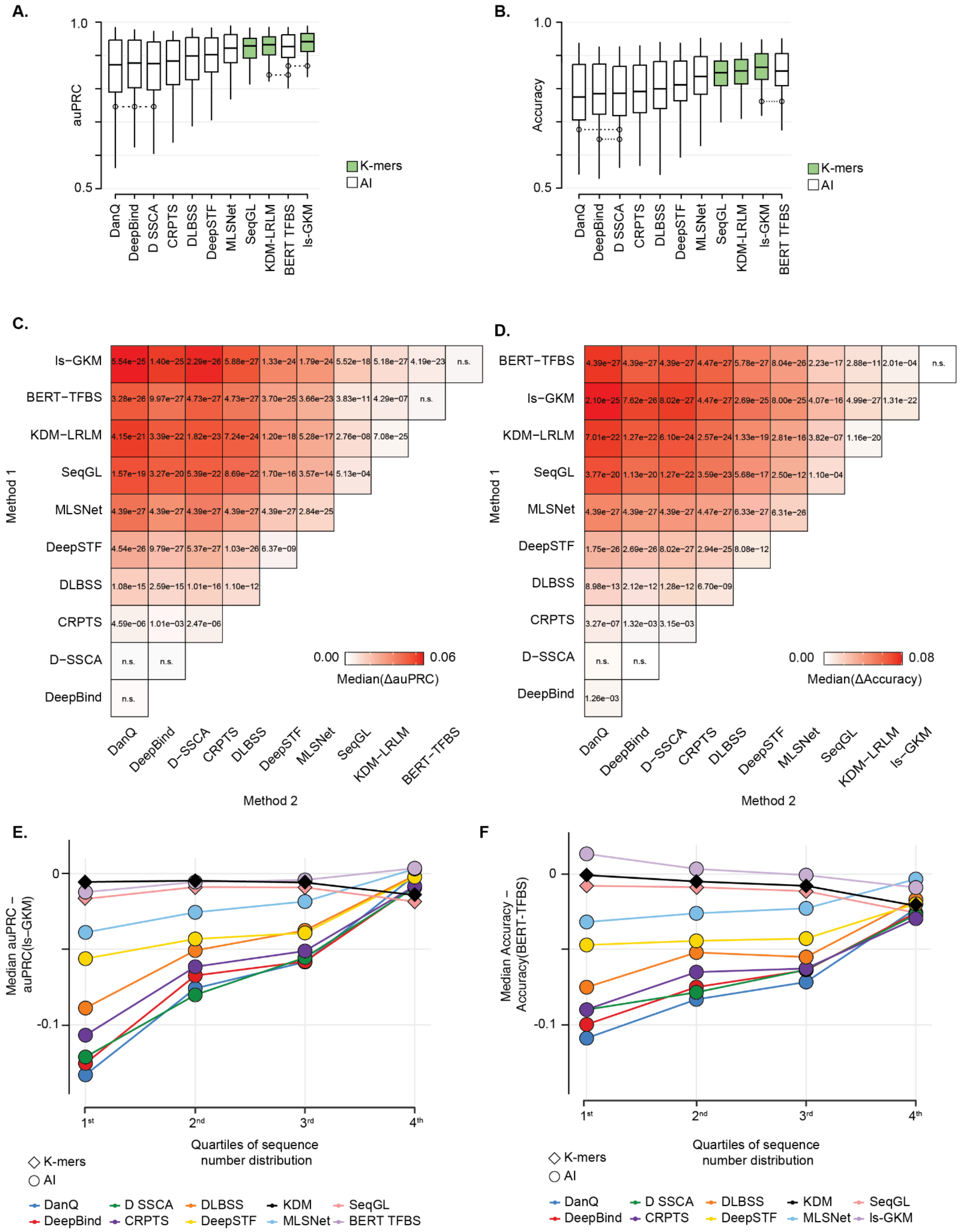


Companion to **Fig. 6D,G**, with auPRC and Accuracy metrics across KDM-LRLM and ten comparator methods (ls-GKM, SeqGL, BERT-TFBS, DanQ, DeepBind, DeepSTF, MLSNet, DLBSS, D-SSCA, CRPTS). **A**. auPRC distributions across TF-benchmark experiments. Green: k-mer–based; white: AI-driven. Boxplots connected by dashed lines indicate distributions not significantly different (paired one-sided Wilcoxon, $p > 0.05$). **B**. Accuracy distributions for the same methods. **C**. Matrix of pairwise median auPRC differences; values are p-values corrected for multiple testing (bonferroni), "n.s." denotes p > 0.05. **D**. As in C for Accuracy. **E**. Median ΔauPRC of each method relative to ls-GKM, with experiments grouped by training-set size quartiles (Q1-Q4). Diamonds: k-mer–based; circles: AI-driven. **F**. Median ΔAccuracy of each method relative to BERT-TFBS, stratified as in E.

#### Figure S6. auPRC and accuracy on RBP-ENCODE.

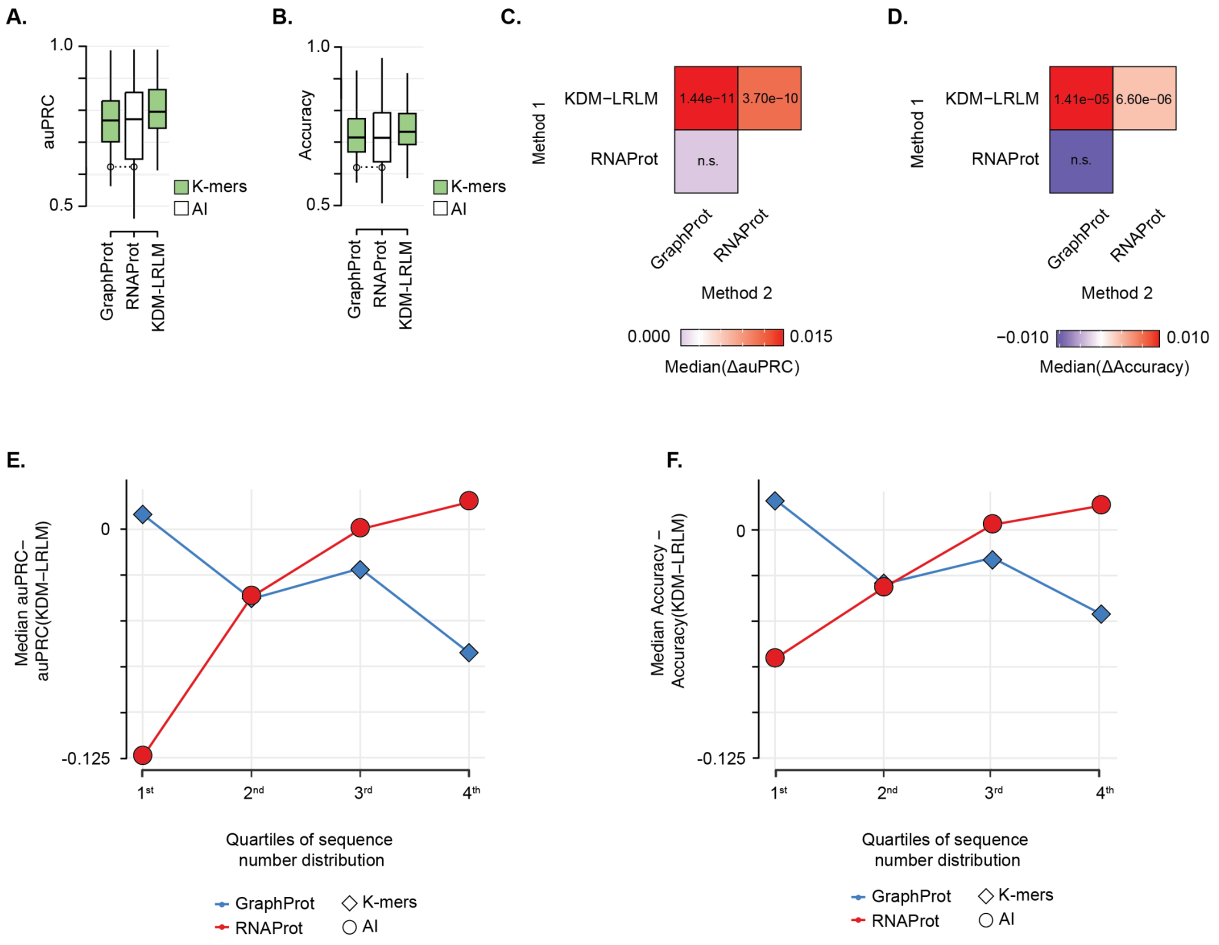


Companion to **Fig. 6E,H**, with auPRC and Accuracy metrics. **A**. auPRC distributions across RBP-ENCODE experiments for KDM-LRLM, GraphProt (k-mer; green), and RNAProt (AI; white). Boxplots connected by dashed lines indicate distributions not significantly different (paired one-sided Wilcoxon, p > 0.05). **B**. Accuracy distributions for the same comparison. **C**. Pairwise median auPRC differences; values are p-values corrected for multiple testing (bonferroni), "n.s." denotes p > 0.05. **D**. As in C for Accuracy. **E**. Median ΔauPRC of GraphProt and RNAProt relative to KDM-LRLM, with experiments grouped by training-set size quartiles (Q1-Q4). Diamonds: k-mer–based; circles: AI-driven. **F**. As in E for ΔAccuracy.
